# Dietary and body mass reconstruction of the Miocene neotropical bat *Notonycteris magdalenensis* (Phyllostomidae) from La Venta, Colombia

**DOI:** 10.1101/2020.12.09.418491

**Authors:** Camilo López-Aguirre, Nicholas J Czaplewski, Andrés Link, Masanaru Takai, Suzanne J Hand

## Abstract

The middle Miocene La Venta bat fauna is the most diverse bat palaeocommunity in South America, with at least 14 species recorded. They include the oldest plant-visiting bat in the New World, and some of the earliest representatives of the extant families Phyllostomidae, Thyropteridae and Noctilionidae. La Venta’s *Notonycteris magdalenensis* is an extinct member of the subfamily Phyllostominae, a group of modern Neotropical animalivorous and omnivorous bats, and is commonly included in studies of the evolution of Neotropical bats, but aspects of its biology remain unclear. In this study, we used a multivariate dental topography analysis (DTA) to reconstruct the likely diet of *N. magdalenensis* by quantitatively comparing measures of molar complexity with that of 25 modern phyllostomid and noctilionid species representing all major dietary habits in bats. We found clear differences in molar complexity between dietary guilds, indicating that DTA is potentially an informative tool to study bat ecomorphology. Our results suggest *N. magdalenensis* was probably an omnivore or insectivore, rather than a carnivore like its modern relatives *Chrotopterus auritus* and *Vampryum spectrum*. Also, we reconstructed the body mass of *N. magdalenensis* to be ∼50 g, which is larger than most insectivorous bats, but smaller than most carnivorous bats. Our results confirm that *Notonycteris magdalenensis* was probably not a specialised carnivore. It remains to be demonstrated that the specialised carnivory ecological niche was occupied by the same lineage of phyllostomines from at least the middle Miocene. Combining our diet and body mass reconstructions, we suggest that *N. magdalenensis* exhibits morphological pre-adaptations crucial for the evolution of specialised carnivory.

## Introduction

With over 200 living species, Phyllostomidae is the family with the highest ecomorphological diversity in the order Chiroptera. The range of dietary niches that phyllostomid bats occupy is unparalleled across Mammalia (Aguirre et al. 2003, Arbour et al. 2019, Dumont et al. 2012, Fleming 1986, Santana et al. 2012). Studies have classified noctilionoid bats into seven dietary guilds: carnivory, frugivory, insectivory, nectarivory, omnivory, piscivory and sanguivory, with insectivory being the most common (Denzinger and Schnitzler 2013, Dumont et al. 2012, Norberg and Rayner 1987, Santana et al. 2012). Reconstructions of dietary evolution in Phyllostomidae suggest that carnivory and nectarivory evolved independently multiple times in different lineages, diverging from an insectivorous common ancestor (Datzmann et al. 2010, Davies et al. 2020, Rojas et al. 2012, Santana and Cheung 2016). Morphological variability associated with diet-based functional demands is thought to have facilitated the ecological radiation and taxonomic diversification in Phyllostomidae (Arbour et al. 2019, Hedrick et al. 2019, Morales et al. 2019, Rossoni et al. 2019, Santana et al. 2012, Shi and Rabosky 2015; Giannini et al. 2020). Traditionally, most studies on dietary ecomorphology in bats have focused on skull morphology to study the form-function link, providing crucial information to understand the range of morphological specialisations (Aguirre et al. 2002, Arbour et al. 2019, Monteiro and Nogueira 2011, Rossoni et al. 2019, Santana and Cheung 2016, Santana and Portugal 2016). Some studies have also provided insights on the role of diet in the diversification of the postcranial skeleton (Gaudioso et al. 2020, Louzada et al. 2019, Norberg and Rayner 1987, Vaughan 1959) and external sensory organs (Brokaw and Smotherman 2020, Leiser-Miller and Santana 2020).

Despite the increasing evidence in support of diet as a key driver of bat evolution, reconstructing the macroevolutionary trajectories of dietary specialisations in bats has been limited by the fossil record (Teeling et al. 2005). Compared to other tetrapod groups, the fossil record of Chiroptera has one of the lowest levels of taxonomic diversity and skeletal preservation, with an estimated 80% of the record missing (Brown et al. 2019, Eiting and Gunnell 2009). Geographic patterns in the completeness of the bat fossil record indicate that the southern hemisphere and Asia are especially underrepresented, obscuring important spatiotemporal information (Brown et al. 2019). In the southern hemisphere, two fossiliferous localities stand out in terms of bat fossil diversity; Riversleigh from the Oligocene to Pleistocene (ca. 25-2 Mya) of Australia with over 40 species (Hand and Archer 2005), and La Venta from the middle Miocene (ca. 12-13 Mya) of Colombia with at least 14 species (Czaplewski et al. 2003).

The La Venta fossil fauna, recovered from the Villavieja Formation in Colombia’s Huila Department, is the richest Cenozoic vertebrate fossil community of northern South America (83 fossil mammal species) (Kay & Madden 1997, Croft 2016). The deposits span a poorly-represented period in the middle Miocene and its fossil mammals have helped define the Laventan South American Land Mammal Age (SALMA) (Madden et al. 1997). The La Venta bat community includes representatives of six families, including extinct representatives of the family Phyllostomidae (*Notonycteris magdalenensis, N. sucharadeus, Palynephyllum antimaster*), and the oldest evidence of modern species of families Noctilionidae (*Noctilio albiventris)* and Thyropteridae (*Thyroptera lavali*) (Czaplewski 1997, Czaplewski et al. 2003). *P. antimaster* represents the earliest phytophagous bat in the New World, a dietary strategy providing a key evolutionary innovation for the order (Czaplewski et al. 2003, Yohe et al. 2015).

La Venta’s *Notonycteris magdalenensis* is known from a dentary fragment containing a complete m1 and the anterior portion of m2, as well as several other isolated teeth, and postcranial remains referred to this species include distal and proximal humeral fragments (Savage 1951, Czaplewski 1997, Czaplewski et al. 2003). When Savage (1951) described the species, he placed *N. magdalenensis* in the phyllostomid subfamily Phyllostominae, a group of Neotropical animalivorous and omnivorous bats. Subsequent phylogenetic analyses have upheld that placement and grouped it consistently in a clade with *Vampyrum spectrum* and *Chrotopterus auritus*, both large-bodied carnivorous species (Czaplewski et al. 2003, Dávalos et al. 2014). Comparisons of dental features and body size indicate that *N. magdalenensis* was larger than *C. auritus* but smaller than *V. spectrum*, the largest bat in the New World. This fossil species is often included in phylogenetic analyses, as a calibration point for dated phylogenies of the bat superfamily Noctilionoidea (Dávalos et al. 2014, Hand et al. 2018, Rojas et al. 2012). It also features in studies focused on the ecological history of phyllostomids, including the adoption of specialised ecological niches such as carnivory by the Middle Miocene (e.g., Baker et al. 2012, Yohe et al. 2015, Simmons et al. 2020). Yet, despite its importance for the study of the evolution of Neotropical bats, further studies focused on *N. magdalenensis* are needed in order to reconstruct its biology.

Mammal teeth show highly specialised structures that vary in relation not only to phylogeny but also to dietary differences, providing informative and widely used proxies to identify niche partitioning and to reconstruct the diets of extinct taxa, including bats (e.g., Czaplewski et al. 2003, Hand et al. 2016, Self 2015, Simmons et al. 2020, Simmons et al. 2008). Studies of the dentition of *N. magdalenensis* by Savage (1951) and Czaplewski et al. (2003) concluded that dental features seen in *N. magdalenensis* that are typically observed in carnivorous bats (Freeman 1988, 1998) were not as highly specialised as those in its living relatives *V. spectrum* and *C. auritus*, suggesting that *N. magdalenensis* probably had a less carnivorous diet than its modern relatives. These dental features in *N. magdalenensis*, compared to its living carnivorous relatives, include relatively lower crown height, more robust crests and cusps, less reduced talonids in lower molars, and less obliquely oriented ectoloph crests, and shorter postmetacrista in upper molars.

Dental topography analysis (DTA) is an informative quantitative approach to study dental morphology within an ecological context and with the potential to be incorporated into modern phylogenetic comparative methods (Allen et al. 2015, Bunn and Ungar 2009, Cooke 2011, Evans et al. 2007, López-Torres et al. 2018, Pineda-Munoz et al. 2017, Prufrock et al. 2016a, Selig et al. 2020). A battery of DTA metrics have been developed in recent decades (Berthaume et al. 2019a, Boyer 2008, Bunn et al. 2011, Evans et al. 2007), further developing the capacity to apply multivariate analyses (Pineda-Munoz et al. 2017). DTA has been widely used to study dietary adaptations in mammals (Lazzari et al. 2008, Selig et al. 2019, Selig et al. 2020), especially in primates (Allen et al. 2015, Berthaume et al. 2020, Bunn and Ungar 2009, Ledogar et al. 2013, López-Torres et al. 2018, Ungar 2004, Ungar et al. 2018, Winchester et al. 2014), and to analyse morphofunctional specialisations (Bunn and Ungar 2009, Bunn et al. 2011), elucidate macroevolutionary trajectories (López-Torres et al. 2018), resolve systematic arrangements (Selig et al. 2020) and reconstruct ecological interactions (Prufrock et al. 2016a) across mammal taxa. In bats, Santana et al. (2011) used DTA to explore diet-based adaptations in the occlusal surface of molars in Phyllostomidae, correlating plant-based diets with higher dental complexity.

We used 3D computational modelling, multivariate DTA and phylogenetic comparative methods to reconstruct the diet of *N. magdalenensis* by comparing the dental complexity of the first lower molar of *N. magdalenensis* with 25 modern phyllostomid and noctilionid species, covering all major dietary categories. We examined three measures of dental complexity (Pineda-Munoz et al. 2017): Dirichlet Normal Energy (DNE), Relief Index (RFI), and Orientated Patch Count Rotated (OPCR). We also reconstructed the body mass (BM) of *N. magdalenensis* using equations developed for Chiroptera based on body size measurements (Gunnell et al. 2009). Based on the phylogenetic relationships of *N. magdalenensis* and previous anatomical comparisons, we hypothesise that it was a large-bodied species, less specialised for carnivory than modern carnivorous phyllostomids. We predict sampled carnivores and *N. magdalenensis* to have high DNE and OPCR values and low RFI values, following general trends found in other mammals. Based on previous comparisons of body proportions with modern relatives, we predict *N. magdalenensis*’ BM to be intermediate between those of *C. auritus* (∼70 g, Vleut et al. 2019) and *V. spectrum* (∼150 g, Amador et al. 2019).

## Methods

The cast of a partial lower left jaw of *N. madgalenensis* with a complete first molar from the collection of the University of California Museum of Paleontology (UCMP39962) was scanned at the University of New South Wales using a U-CT (Milabs, Utrecht) with 55kV and 0.17 mA, ultrafocused setting at a resolution of 30-50 μm (Fig. 1). Our comparative sample consisted of 25 modern noctilionoid bat species of known diet from two noctilionoid bat families (Phyllostomidae and Noctilionidae; supplementary table 1). Together these species represent all major dietary guilds (i.e., carnivory, frugivory, insectivory, nectarivory, omnivory, piscivory and sanguivory) recognised in bats (Table 1 and Fig. 1). For each species, a single lower left first molar (m1) from adult specimens that did not show excessive wear was analysed, in order to avoid methodological artifacts in our DTA (Pampush et al. 2016a). 3D models of modern species were retrieved from Shi & Rabosky (2018) based on μCT scans available on MorphoSource, except for an additional 3D model of a lower mandible of *V. spectrum* provided by S. Santana upon request. Individual m1s were segmented from adjacent teeth and bone using MIMICS v. 20 software (Materialise NV, Leuven, Belgium). Additionally, following standard protocols for DTA, individual m1s were cropped below the cingulid to isolate the crown (Prufrock et al. 2016a, Ungar et al. 2018). Dávalos et al.’s (2014) phylogeny of the Phyllostomidae was pruned to match our sample and used as phylogenetic scaffold for phylogenetic comparative analyses.

**Table 1.** Description of dietary guild classification used in this study.

| Dietary Guild | Description |
| --- | --- |
| Carnivory | Diet including or dominated by the consumption of terrestrial vertebrates |
| Frugivory | Diet dominated by the consumption of fruits |
| Insectivory | Diet dominated by the consumption of invertebrates |
| Nectarivory | Diet dominated by the consumption of pollen |
| Omnivory | Diet composed of a mix of fruit, pollen and invertebrates |
| Piscivory | Diet dominated by the consumption of fish |
| Sanguivory | Diet dominated by the consumption of blood |

**Figure 1.**
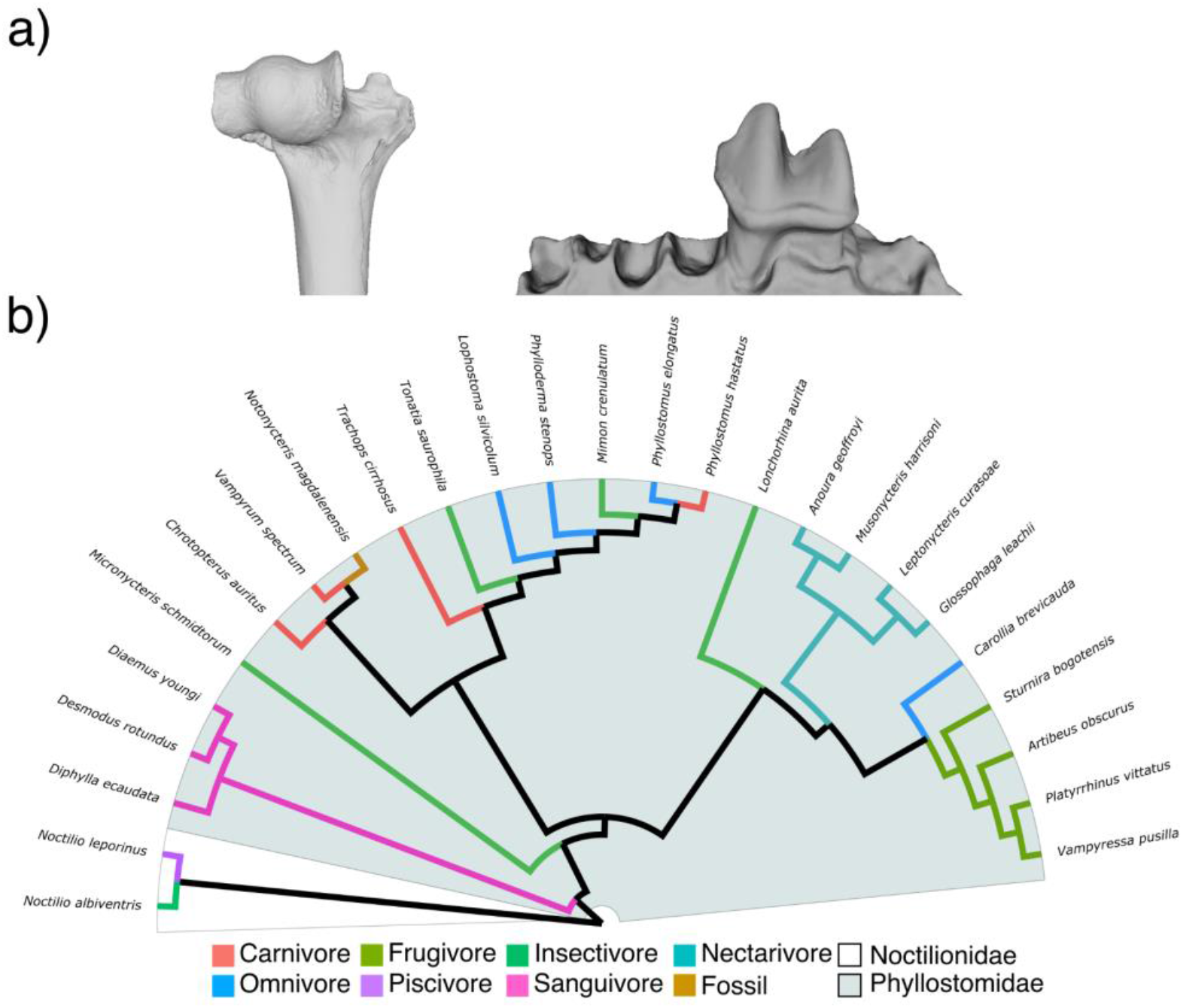
Fossil and modern species analysed in this study. a) Partial humerus (UCMP 38990; left) and lower left jaw (UCMP 39962; right) of *N. magdalenensis* used in this study for body mass (BM) reconstruction and dental topography analysis (DTA); b) Evolutionary relationships of species analysed in this study, modified from Dávalos et al. (2014). Branch colours represent dietary guilds and the fossil taxon.

### Body mass estimate

Based on a dataset of 1160 extant bats, Gunnell et al. (2009) demonstrated that regression analysis of skeletal elements can be an accurate proxy to infer body mass (BM) in extant and extinct bats, with humeral shaft diameter and first upper/lower molar area providing best estimates. We used lower first molar area and humeral shaft diameter to reconstruct the BM of *N. magdalenensis* (Gunnell et al. 2009, Hand et al. 2016, Hand et al. 2018, Hand et al. 2015a, Jones et al. 2018). Lower first molar area was calculated from a 3D model of a complete m1 extracted from a fragmented left lower jaw (UCMP 39962). Only fragmented humeral elements have been reported for *N. magdalenensis*, limiting our capacity to obtain measures of mid-shaft diameter. Humeral dorsoventral diameter of the shaft at the distal end of the deltopectoral ridge of a proximal humerus fragment (UCMP 38990) was retrieved from Savage (1951), and proximal-most shaft diameter of the humerus was extracted from the 3D model of the distal end of a partial humerus (UCMP 38990; Fig 1). A cast of UCMP 38990 was scanned at the University of New South Wales following the same protocol as described above. The BM estimate based on m1 size was used to compare *N. magdalenensis* with bats from modern dietary guilds. BM of modern bat species was retrieved from the literature (Amador et al. 2019, Curtis and Santana 2018, Gunnell et al. 2009, Hand et al. 2018, Jones et al. 2018, Molinari et al. 2017).

### Dental topographic analysis

Segmentation and processing practices (e.g. cropping and smoothing) of 3D meshes can impact features of the models (e.g. triangle count and mesh resolution) that are determinant for DTA (Berthaume et al. 2019b). Hence, all meshes were standardised following recommendations for DTA; 3D models were simplified to 10,000 faces and smoothed using 30 iterations with lambda set at 0.6 (Berthaume et al. 2019b, Pampush et al. 2016b). After standardisation of meshes, three widely used dental topography metrics were analysed (Fig. 2): DNE (Bunn et al. 2011), OPCR (Evans et al. 2007), and RFI (Boyer 2008). DNE is a measure of the degree of curvature of the crown (Fig. 2a); this correlates with shearing capacity, and is usually higher in animalivores (Allen et al. 2015, Ledogar et al. 2018, Ledogar et al. 2013, López-Torres et al. 2018, Ungar et al. 2018). OPCR is an estimate of surface complexity of the crown based on the number of patches facing in the same direction (Evans et al. 2007, López-Torres et al. 2018, Pineda-Munoz et al. 2017, Selig et al. 2020). As for DNE, species with a higher OPCR have more complex tooth surfaces (Fig. 2b), usually associated with more biomechanically demanding diets such as folivory and animalivory (Cooke 2011, Evans et al. 2007, Santana et al. 2011). RFI is a ratio of the three-dimensional and two-dimensional area of the tooth crown (Fig. 2c), and provides a measure of crown height and effectively hypsodonty (Allen et al. 2015, Boyer 2008, Cooke 2011); low RFI values reflect flat crown surfaces commonly found in mammalian frugivores, whereas high values represent tall cusps such as those found in granivores and insectivores. Incorporating several metrics of dental complexity to inform dietary reconstructions has been suggested to provide more robust characterisations than studies based on a single metric (Pineda-Munoz et al. 2017). All DTA were performed with the R package molaR (Pampush et al. 2016b).

**Figure 2.**
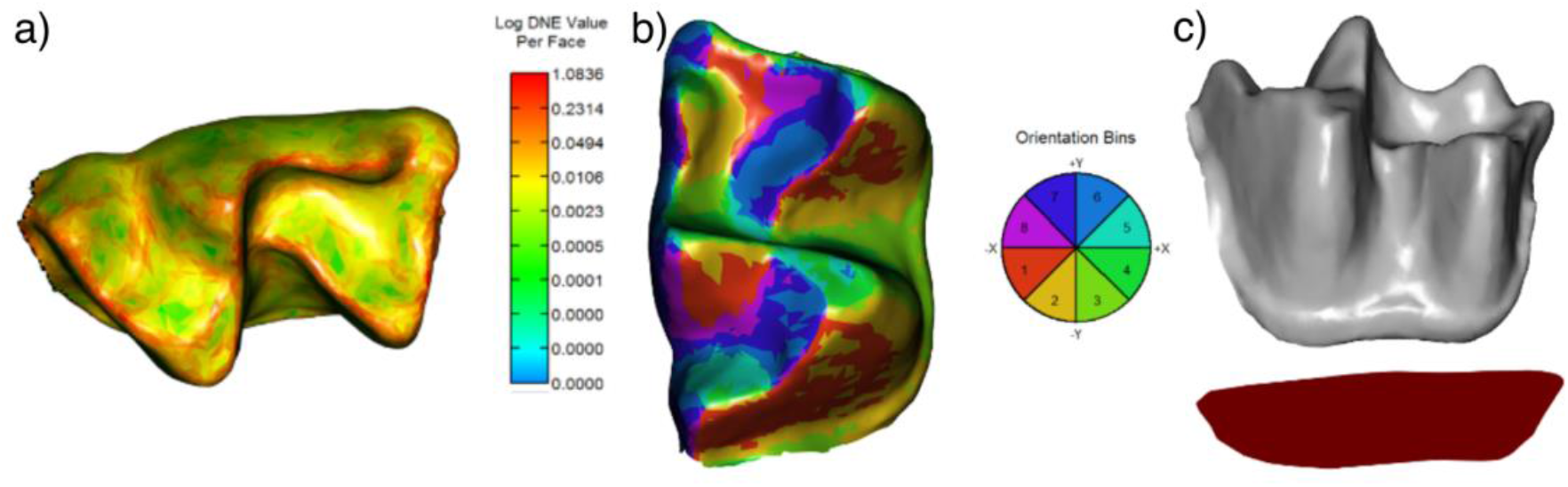
Reconstructed meshes showing topographic maps of DNE (a), OPCR (b) and RFI (c) for the lower m1 of *N. madgalenensis* (UCMP 39962).

### Dietary reconstruction

Boxplots were used to visually compare values of DNE, OPCR and RFI in *N. magdalenensis* with those of modern bats of known dietary guilds. A one-way ANOVA was used to test for differences in DNE, OPCR and RFI across our groups. Presence of phylogenetic structuring in our dental complexity data was tested using a multivariate Kmult (K−) statistic implemented in the “physignal” function in *Geomorph* 3.2.1 (Adams 2014), using a pruned version of Dávalos et al.’s (2014) phylogeny. Given the strong phylogenetic signal in our dataset, a Principal Component Analysis (PCA herein morphospace) and a phylogenetic PCA (pPCA herein phylogeny-corrected morphospace) were performed to reduce the dimensionality of our dental complexity data and visualise patterns of variation, using the “phylomorphospace” and “phyl.pca” functions of the R package phytools (Revell 2012). A morphospace plot allowed us to elucidate general patterns of differentiation across dietary guilds, whereas the phylogeny-corrected morphospace allowed us to discern the influence of phylogenetic kinship in dental complexity variation. A Linear Discriminant Analysis (LDA) was used to classify the diet of *N. magdalenensis* as one of seven dietary guilds, using the “lda” function in the R package MASS. Differences in phylogenetic kinship and patterns of morphological diversity were further explored with a tanglegram in order to: 1) identify dental complexity similarities between *N. magdalenensis* and other bats of known dietary guilds and 2) detect instances of morphological innovation and convergence across our sample. Loadings of the first two principal components (PC) of the pPCA (explaining > 90% of phenotypic variation) were used as phylogeny-corrected phenotypic data. First, the phenotypic dendrogram was built with a hierarchical clustering analysis of the PC loadings, using the “upgma” function in the R package phangorn (Schliep 2011). Next, we compared the position of each species across the phylogeny and morphological dendrogram using the tanglegram produced by the “cophylo” function in the R package phytools (Revell 2012). Parallel lines linking a species across the phylogeny and dendrogram indicate similarities, whereas diagonal links indicate a phylogenetic-phenotypic mismatch.

## Results

### Body mass estimate

Depending on the proxy, BM estimates of *N. magdalenensis* ranged from 53.05 g (m1 area), 103.39 g (humeral shaft diameter) to 133.35 g (humeral shaft diameter from Savage 1951). However, a direct measure of humeral diameter at the mid-shaft could not be made (this point is not preserved in fossil material), and this is generally the narrowest point of the bat humerus. Hence, our BM estimates for *N. magdalenensis* based on humeral shaft diameter potentially overestimate body mass and were therefore not included in further analyses. All further analyses and discussion are based on the estimate derived from m1 area (∼50 g). Compared to modern species in our sample, the body mass of *N. magdalenensis* was greater than the average for frugivores, insectivores, nectarivores, omnivores and sanguivores, was similar to the average for piscivores, and smaller than most of the carnivores (Fig. 3). *N. magdalenensis* was smaller than both of its modern relatives (*C. auritus* BM = 80 g, *V. spectrum* BM = 170 g).

**Figure 3.**
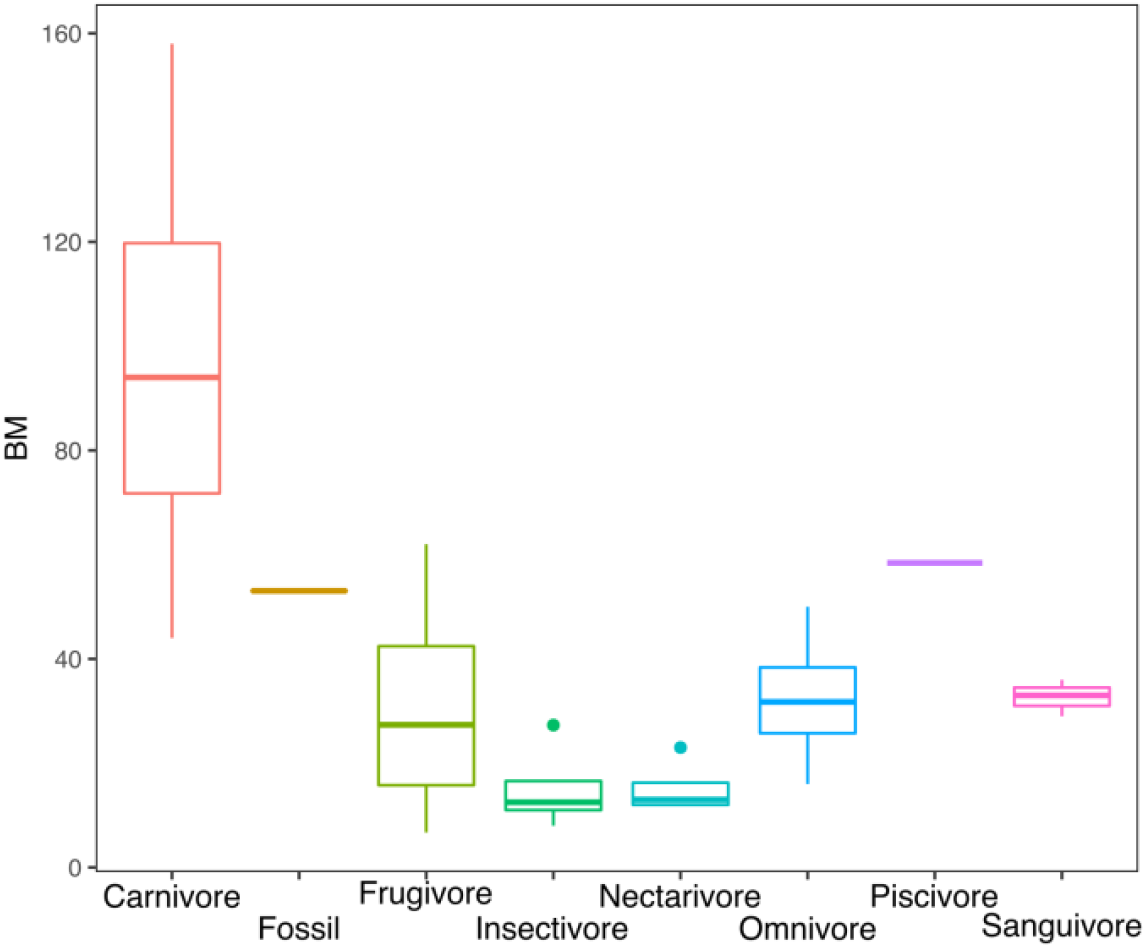
Boxplots of BM distributions of modern taxa pooled by dietary guilds and BM reconstruction of *N. magdalenensis* based on m1 area.

### Dietary differences in dental complexity

In our sample, nectarivores and sanguivores consistently had the lowest values of DNE and OPCR, whereas frugivores and sanguivores had the lowest values of RFI (Fig. 4). Omnivores and piscivores had the highest DNE values, followed by frugivores and carnivores (Fig. 4a). Frugivores had the highest OPCR values, followed by carnivores, insectivores, omnivores and piscivores with very similar values (Fig. 4b). Piscivores and sanguivores had the highest RFI values, followed by carnivores, whereas insectivores and omnivores showed intermediate values (Fig. 4c). Across the three metrics, *N. magdalenensis* overlapped with values for insectivores and omnivores and showed lower values than its carnivorous close relatives (Fig. 4). One-way ANOVA revealed statistically significant differences in dental complexity metrics across dietary guilds (F = 2.46, *p* = 0.05), and the K-statistic revealed a significant phylogenetic signal in our data (K = 0.686, Z = 2.273, *p* = 0.006).

**Figure 4.**
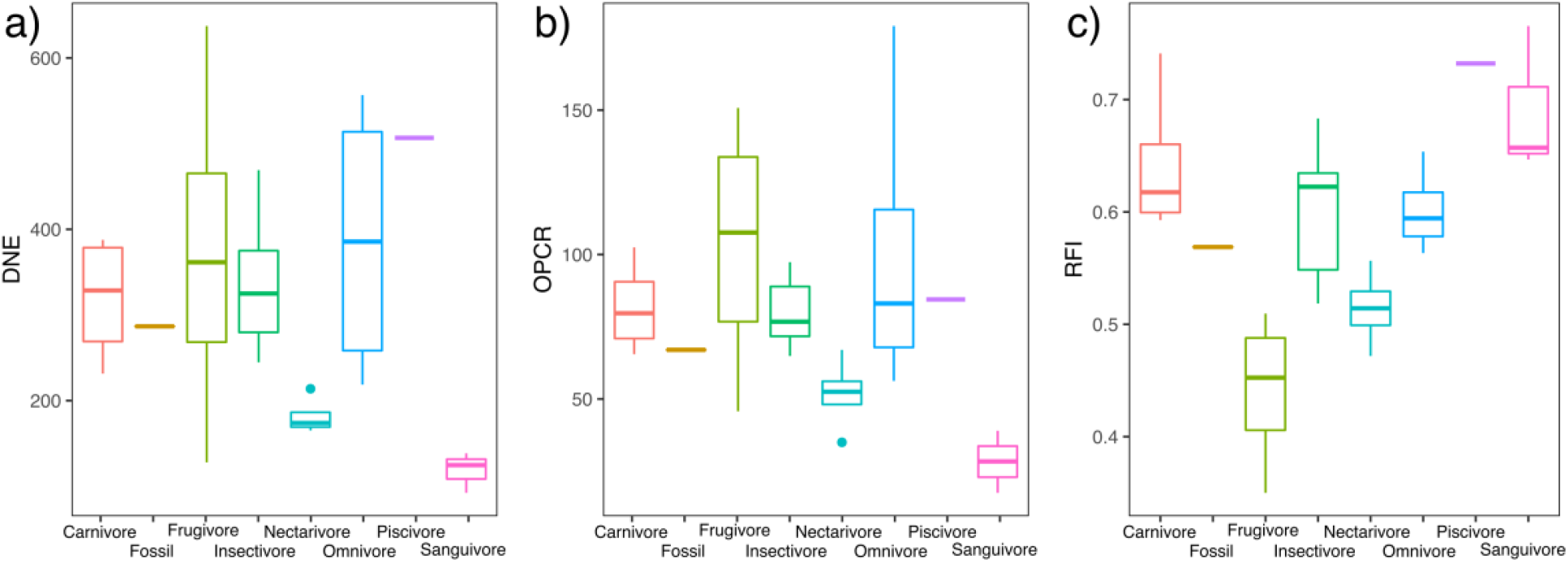
Boxplots of DTA results for DNE (a), OPCR (b) and RFI (c) of modern taxa pooled by dietary guilds and *N. magdalenensis*.

Dietary groups showed similar patterns of phenotypic variation across morphospace and phylogeny-corrected morphospace (Fig. 5). In morphospace (first two PCs explained 95.77% of variation), frugivores, nectarivores and sanguivores occupied non-overlapping regions, whereas carnivores, insectivores and omnivores overlapped, each showing different levels of dispersion across morphospace (Fig. 5a). Morphospace also indicates phylogenetic structuring in morphological variation and dietary specialisations, with desmodontines (all sanguivores), glossophagines (all nectarivores) and stenodermatines (all frugivores) occupying exclusive areas in morphospace. Species showed a similar arrangement in phylogeny-corrected morphospace (first two PCs explained 95.59% of variation), indicating that despite the effect of evolutionary relatedness, dental variation exhibits ecology-based patterns of variation (Fig. 5b). Frugivores, nectarivores and sanguivores occupied unique subspaces, and carnivores, insectivores and omnivores overlapped around the origin. The specialised piscivore *Noctilio leporinus* always separated from all other guilds but the omnivore *N. albiventris* overlapped in morphospace with other animalivore guilds. Across morphospaces *N. magdalenensis* was always nested within the insectivore subspace, close to the omnivore subspace and outside of the carnivore subspace.

**Figure 5.**
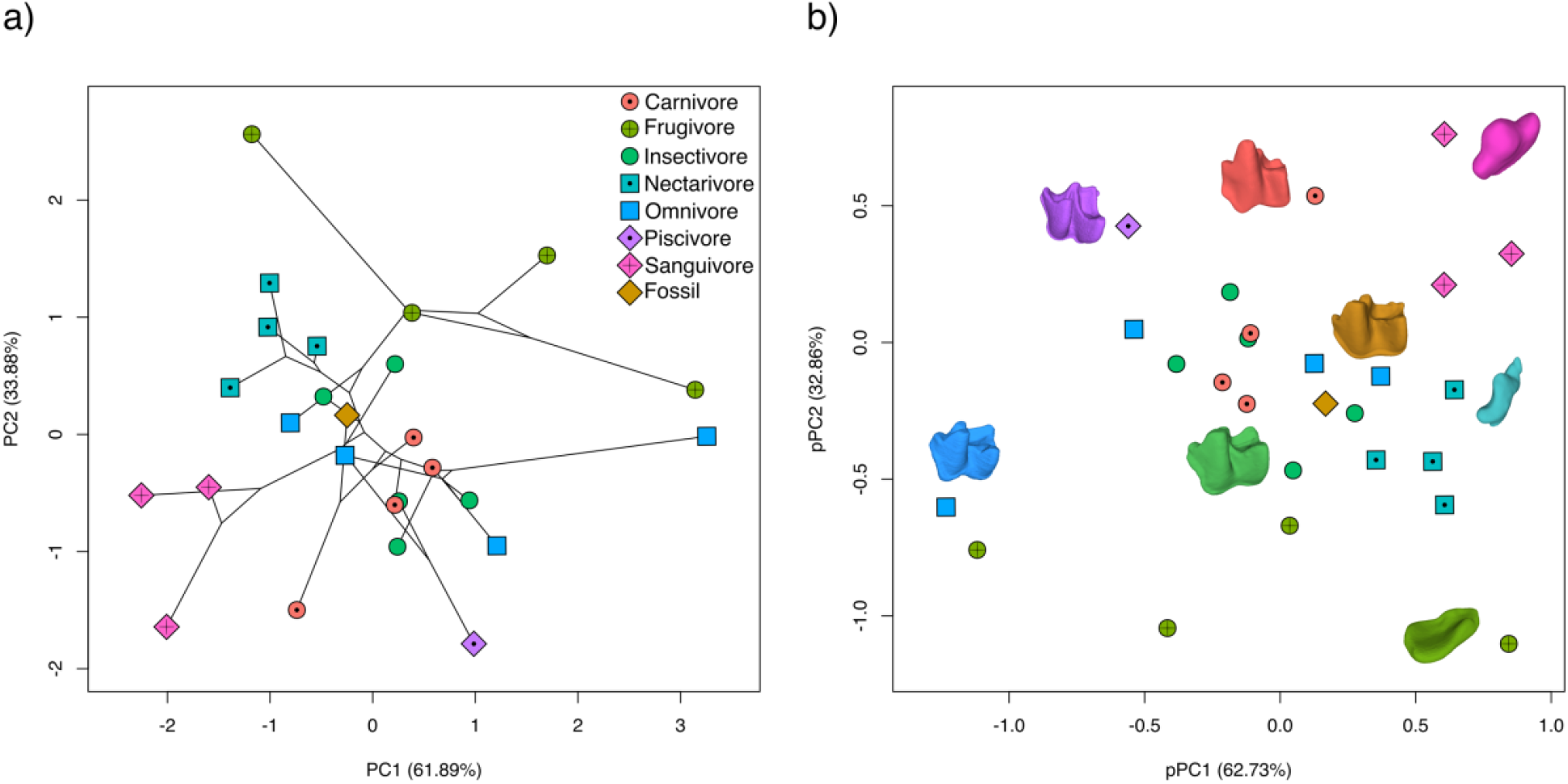
Morphospace (PCA, a) and phylogeny-corrected morphospace (pPCA, b) based on DTA (DNE, OPCR and RFI). Dot colours represent dietary guild categories (carnivory, frugivory, insectivory, nectarivory, omnivory, piscivory, sanguivory) and fossil species (*N. magdalenensis*). 3D models of m1 illustrate dental diversity in sampled taxa, colours matching dietary and fossil categories: Carnivory (*Phyllostomus hastatus*), frugivory (*Sturnira bogotensis*), insectivory (*Micronycteris schmidtorum*), nectarivory (*Musonycteris harrisoni*), omnivory (*Lophostoma silvicolum*), piscivory (*Noctilio leporinus*), sanguivory (*Desmodus rotundus*), and fossil (*Notonycteris magdalenensis*).

### Dietary reconstruction

Tanglegram showed a similar placement of the piscivore, two carnivores and one insectivore species in the phylogeny and dendrogram, and a consistent clustering of sanguivore and nectarivore taxa (Fig. 6). A mismatch between the placement of *N. magdalenensis* in the phylogeny and phenotypic dendrogram was also evident. The phenotypic dendrogram clustered *N. magdalenensis* with the insectivorous *Lonchorhina aurita*, and this cluster with the omnivorous *Phylloderma stenops* in our sample.

**Figure 6.**
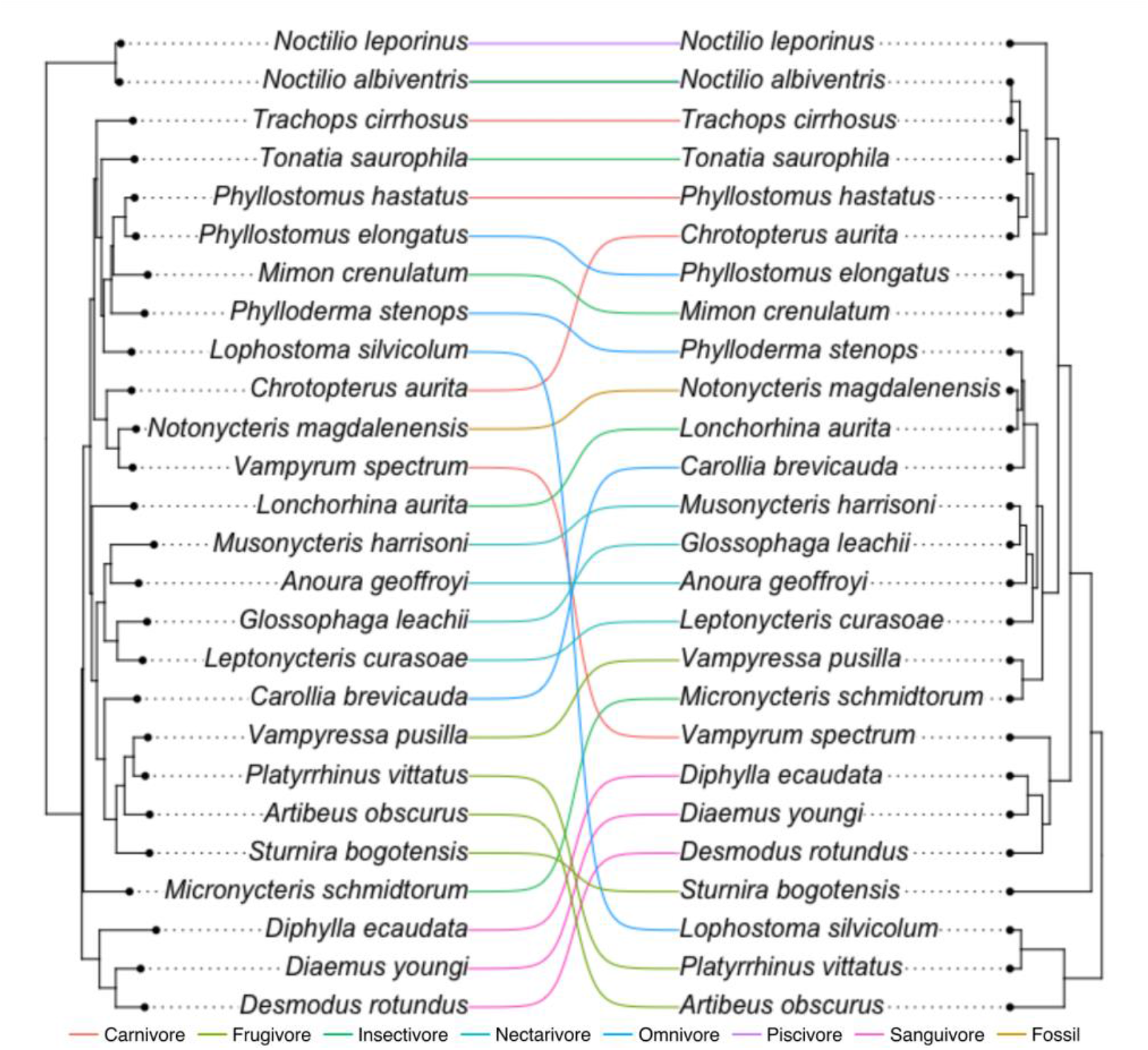
Tanglegram of evolutionary relatedness (left) and phenotypic dendrogram (right) of bat dental complexity. Dotted lines indicate the position of the same species in both panels. Colour of dotted lines represents dietary guild categories and fossil taxon.

LDA correctly classified 64% of modern species in our sample into their dietary guild when all three DTA metrics were analysed, and 36% of modern species when DNE and OPCR only were analysed (Table 2). When DNE, OPCR and RFI were combined, 100% of frugivores, nectarivores, piscivores and sanguivores were correctly classified, followed by omnivores (75%) and insectivores (20%). Excluding RFI from the LDA decreased the accurate classification of frugivores by 75%, of omnivores by 50%, of nectarivores by 25% and of insectivores by 20%. Both iterations of LDA classified *N. magdalenensis* as an insectivore-omnivore, with a posterior probability of 25.35% and 30.92% (using all metrics) and 14.36% and 22.03% (excluding RFI). Both iterations of LDA failed to correctly classify all modern carnivores, with 50% classified as omnivores and 25% as insectivore and sanguivore. Nevertheless, carnivory had the second highest posterior probability (> 20%) across all modern carnivores (supplementary table 2). In contrast, carnivory had the fourth highest LDA posterior probability (17.6%) when classifying *N. magdalenensis*’s diet.

**Table 2.** Predicted dietary guild classification based on LDAs analysing DNE, OPCR and RFI (ALL), and excluding RFI (DNE+OPCR).

| Species | Dietary category | Predicted ALL | Predicted DNE+OPCR |
| --- | --- | --- | --- |
| *Anoura geoffroyi* | Nectarivore | Nectarivore | Nectarivore |
| *Artibeus obscurus* | Frugivore | Frugivore | Frugivore |
| *Carollia brevicauda* | Omnivore | Omnivore | Nectarivore |
| *Chrotopterus aurita* | Carnivore | Insectivore | Frugivore |
| *Desmodus rotundus* | Sanguivore | Sanguivore | Sanguivore |
| *Diaemus youngi* | Sanguivore | Sanguivore | Sanguivore |
| *Diphylla ecaudata* | Sanguivore | Sanguivore | Sanguivore |
| *Glossophaga leachii* | Nectarivore | Nectarivore | Nectarivore |
| *Leptonycteris curasoae* | Nectarivore | Nectarivore | Sanguivore |
| *Lonchorhina aurita* | Insectivore | Nectarivore | Nectarivore |
| *Lophostoma silvicolum* | Insectivore | Insectivore | Frugivore |
| *Micronycteris schmidtorum* | Insectivore | Nectarivore | Omnivore |
| *Mimon crenulatum* | Insectivore | Omnivore | Piscivore |
| *Musonycteris harrisoni* | Nectarivore | Nectarivore | Nectarivore |
| *Noctilio albiventris* | Omnivore | Omnivore | Omnivore |
| *Noctilio leporinus* | Piscivore | Piscivore | Piscivore |
| *Phylloderma stenops* | Omnivore | Carnivore | Carnivore |
| *Phyllostomus elongatus* | Omnivore | Omnivore | Piscivore |
| *Phyllostomus hastatus* | Carnivore | Omnivore | Omnivore |
| *Platyrrhinus vittatus* | Frugivore | Frugivore | Insectivore |
| *Sturnira bogotensis* | Frugivore | Frugivore | Nectarivore |
| *Tonatia saurophila* | Insectivore | Carnivore | Frugivore |
| *Trachops cirrhosus* | Carnivore | Omnivore | Omnivore |
| *Vampyressa pusilla* | Frugivore | Frugivore | Carnivore |
| *Vampyrum spectrum* | Carnivore | Sanguivore | Nectarivore |
| *Notonycteris magdalenensis* |  | Omnivore | Insectivore |

## Discussion

DTA reconstructed the diet of *N. magdalenensis* as an omnivorous or insectivorous species, and distinct from its modern sister taxa *C. auritus* and *V. spectrum*, both of which are carnivores. Body mass estimates suggest that *N. magdalenensis* was a large-bodied bat (∼50 g), larger than most modern phyllostomids and bats in general (Gunnell et al. 2009), but smaller than its modern sister taxa. The reconstructed diet and BM of *N. magdalenensis* suggest this species could represent a transitional stage between small-bodied ancestral insectivorous phyllostomids (Simmons et al. 2020) and large, specialised modern carnivores, illustrating the importance of body size increases for the evolution of carnivory in Chiroptera (Santana and Cheung 2016; Giannini et al. 2020). DTA offers an additional tool to help reconstruct and differentiate dietary niches in bats.

### DTA and dietary specialisations in bats

Functional demands associated with the biomechanical processing of food has been linked to morphological adaptations of the feeding apparatus and sensory organs in bats (Arbour et al. 2019, Brokaw and Smotherman 2020, Jacobs et al. 2014, Leiser-Miller and Santana 2020, Nogueira et al. 2009, Rossoni et al. 2019, Santana et al. 2010, Santana et al. 2012). Food item hardness has been linked with the robustness of the jaw and the length of the rostrum (Arbour et al. 2019; Freeman 1981, 1998, 2000). Studies have also found evidence suggesting dietary adaptations correlate with postcranial morphological differences, revealing the interaction between diet (i.e. food processing) and locomotion (i.e. foraging strategies) has an overarching effect on the ecomorphology of bats (Gaudioso et al. 2020, Morales et al. 2019, Norberg and Rayner 1987). Scapular morphology in bats has been correlated with the convergent evolution of dietary adaptations (Gaudioso et al. 2020). Dental morphology has been shown to have both a phylogenetic and ecological signal, reflecting evolutionary relatedness as well as ecological differences within Chiroptera (e.g. Freeman 1988, 1995, 2000, Hand 1985, 1996, 1998, Santana et al. 2011, Self 2015, Simmons et al. 2016, Zuercher et al. 2020). Our results also detected a dual phylogenetic and ecological signal in dental complexity variation and highlights DTA as a potential tool in the study of dental ecology and evolution in bats. Santana et al. (2011) used OPCR to assess diet-based ecomorphological differences between noctilionoid bats. By applying DNE, OPCR and RFI, ours is the first study to apply multivariate DTA to investigate bats, fossil and living. There is potential for future studies to explore a variety of ecological and evolutionary questions, including the many cases of convergent diets between different bat clades (e.g. frugivory, nectarivory and carnivory) and the reconstruction of ancestral states to investigate dietary diversification in Chiroptera. Our results of DTA-based LDA for carnivorous bats contrast with previous studies analysing carnivorous crocodylimorphs (Melstrom and Irmis 2019) and non-volant mammals (Pineda-Munoz et al. 2017) that revealed a greater differentiation of this diet. One possible explanation is that our approach was based on a single tooth, due to the nature of the *N. magdalenensis* fossil material, whereas previous studies analysed whole toothrows, which could capture a greater suite of dental specialisations.

Our study focused on reconstructing the likely diet of the extinct phyllostomid *Notonycteris magdalenensis* by comparing its dental morphology with that of modern members of this family. Phyllostomidae is recognised as the greatest adaptive radiation of any mammalian family (Fleming et al. 2020, Monteiro and Nogueira 2011, Rossoni et al. 2017). Adoption of novel dietary niches and morphological innovation have been linked to the diversification of this family (Dumont et al. 2012, Freeman 2000, Hedrick et al. 2019, Rossoni et al. 2019). Representing less than 15% of modern global bat diversity (∼200 species), phyllostomid bats have successfully colonised almost every dietary niche found in the entire chiropteran order, with the exception of piscivory (Fleming et al. 2020, Monteiro and Nogueira 2011, Rossoni et al. 2017). Previous studies have suggested that different phyllostomid clades adapted to nectarivory convergently, reflecting the ecomorphological evolvability in phyllostomids (Datzmann et al. 2010, Rojas et al. 2016). Low DNE and OPCR values found in our sanguivore and nectarivore bats indicate evolutionary convergence of reduced occlusal curvature in species with liquid diets. Loss or reduction of incisors and cheek teeth have been identified as convergent adaptations to liquid diets in Chiroptera (e.g., Freeman 1988, 1998, Berkovitz and Shellis 2018, Bolzan et al. 2015). Nevertheless, sanguivorous and nectarivorous bats also show divergent specialised cranial morphologies adapted to each diet, with these dietary innovations following different evolutionary trajectories (Rossoni et al. 2017, Rossoni et al. 2019). Sanguivorous bats have a short rostrum, enlarged and procumbent upper incisors adapted for slicing, and reduced cheek teeth and lower incisors (Berkovitz and Shellis 2018). Nectarivorous bats exhibit elongated rostra and reduction or loss of incisors and modified cheek teeth (Freeman 1988, 1995). This divergence is also reflected in marked differences in RFI values between sanguivores and nectarivores and non-overlapping regions occupied in morphospace (Fig. 5). High RFI values found in sanguivore bats reflect the reduction of molar width during the evolution of liquid diets in bats.

A previous study of dental complexity in bats found higher OPCR in frugivores compared to insectivores and omnivores (Santana et al. 2011), unlike the pattern recorded in rodents and primates (Evans et al. 2007, López-Torres et al. 2018, Selig et al. 2019, Selig et al. 2020, Winchester et al. 2014). Our results (i.e. high OPCR in frugivores compared to animalivores and omnivores) capture the uniquely complex teeth of frugivorous phyllostomid bats, compared to other frugivorous mammals, including pteropodid bats. Non-phyllostomid frugivorous mammals tend to have flatter occlusal surfaces due to reduced shearing surfaces (e.g. cusps and crests), whereas the lower molars of phyllostomid frugivores retain a labial crest formed by the metaconid and paraconid (Berkovitz and Shellis 2018). Frugivorous species in our sample had the lowest RFI values of any dietary group, showing low-crowned teeth similar to those in other mammal groups (Boyer 2008, Bunn and Ungar 2009, Prufrock et al. 2016b). Carnivorous and piscivorous species occupied different subregions of morphospace, indicating demands of feeding on terrestrial and aquatic vertebrates shaped dental complexity differentially in each group. Cranial morphology of piscivore bats has been found to be significantly different from other animalivorous bats, possibly due to differences in the masticatory and foraging biomechanics between both groups (Santana and Cheung 2016). Future studies could expand to include a wider range of bat species and families to test convergent evolution between more distantly-related taxa.

### Dietary and body mass reconstruction of *N. magdalenensis*

Our inferences of the diet and body mass of *N. magdalenensis* may have bearing on broader understanding of the evolution of carnivory in Phyllostomidae. Some studies have estimated that carnivory had evolved in Phyllostomidae by at least the middle Miocene, based on the dental morphology of *N. magdalenensis* (Czaplewski et al. 2003, Savage 1951, Simmons et al. 2020). Further comparisons of measurements of cranial and postcranial remains have been used to suggest this species was larger than *C. auritus* but smaller than *V. spectrum* (Czaplewski et al. 2003, Savage 1951, Simmons et al. 2020). Giannini et al. (2020) explored the evolution of body mass in extant noctilionoid bats, including all of the phyletic lineages and dietary modes exhibited by them. These authors hypothesised that body-size stability early in the evolution of phyllostomids provided great potential for size increases and decreases among them. They suggest that the increase in BM that enabled specialised carnivory, ultimately resulting in the very large-bodied Vampyrini (*C. auritus* and *V. spectrum*), probably occurred relatively late (late Miocene rather than middle Miocene; molecular divergence dating of Amador et al. 2016), which is after the time of the La Venta fauna and of *N. magdalenensis*. Our results suggest that *N. magdalenensis* may have been a large insectivore or omnivore, with an intermediate BM between insectivores and carnivores in our sample, and are concordant with (or at least, do not refute) a later appearance of specialised carnivorous phyllostomids.

In our DTA study, *N. magdalenensis* had lower values of DNE, OPCR and RFI than the modern carnivores and piscivores in our sample, indicating lower occlusal curvature, dental complexity, and crown height (i.e. lesser capacity to slice vertebrate prey). LDA classified *N. magdalenensis* primarily as an insectivore (posterior probability of 35%), and secondarily as an omnivore (posterior probability of 21.24%), whereas there was only a 16.21% posterior probability of carnivory. Our tanglegram based on the same data also suggests an omnivorous or insectivorous diet for *N. magdalenensis*, which groups within a cluster of omnivorous and insectivorous species, distant from carnivores or its sister taxa. Based on our results, we hypothesise that *N. magdalenensis* represents a transitional form between modern carnivory and the ancestral insectivore state.

Moreover, an omnivorous or insectivorous diet for *N. magdalenensis* also supports the observations by Savage (1951) and Czaplewski et al. (2003). Their qualitative studies of the dentition of *N. magdalenensis* found that dental features in *N. magdalenensis* typically observed in carnivorous bats (Freeman 1988, 1998) were not as highly developed as those same features in its living relatives *V. spectrum* and *C. auritus*, suggesting that *N. magdalenensis* probably had a less carnivorous diet than its modern relatives. These features include relatively lower crown height in *N. magdalenensis*, more robust crests and cusps, less reduced talonids in lower molars, and less obliquely oriented ectoloph crests, and shorter postmetacrista in upper molars (Czaplewski et al. 2003).

With the exception of *Trachops cirrhosus* (∼35 g), all carnivorous phyllostomids have average body masses higher than 70 g, well above the average for any other diet (Moyers Arévalo et al. 2018, Norberg and Fenton 1988, Santana and Cheung 2016). *V. spectrum*, thought to be the closest living relative of *N. magdalenensis*, is the largest bat in the New World (BM ∼170 g), and regarded to be an example of the coevolution of increased body size and carnivory in Phyllostomidae (Arévalo 2020, Moyers Arévalo et al. 2018). Our estimate of BM for *N. magdalenensis* (∼53 g) based on m1 area indicates that this species was smaller than most phyllostomid carnivores, larger than most insectivores, and within the range of omnivorous phyllostomids. In agreement with our DTA, this could indicate that *N. magdalenensis* was not a specialised carnivore and could have instead had an insectivorous or omnivorous diet, similar to other omnivorous, large-bodied Miocene noctilionoid species (Hand et al. 2015b, 2018). Other studies have estimated phyllostomid BM ancestral state range as between 11-12 g, close to the ancestral state for Noctilionoidea (9-12 g) and modern Chiroptera (12-14 g) (Arévalo 2020, Giannini et al. 2012, 2020, Moyers Arévalo et al. 2018). This suggests relative evolutionary stasis in body size during the early evolution of modern bat families, most of which had evolved and/or had appeared in the fossil record before the early Oligocene (∼35 Mya) (Arévalo 2020, Giannini et al. 2012, Moyers Arévalo et al. 2018). Within Phyllostominae, a significant shift to increased size to more than 50 g has been recovered for the clade including *C. auritus* and *V. spectrum* (divergence dated at around 16 Mya), highlighting a rapid gain of > 30 g in the first 10 My following the origin of Phyllostomidae (Arévalo 2020). Based on the age of the La Venta bat fauna (12-13 Mya), *N. magdalenensis* can be interpreted as fossil evidence for this evolutionary increase in body size within Phyllostominae (Arévalo 2020), and we propose it represents a transitional stage of body size between the small insectivore phyllostomid ancestor and the large modern carnivore phyllostomids (Baker et al. 2012, Datzmann et al. 2010). Taking our DTA and BM reconstruction together, we hypothesise that during the evolution of carnivory in Phyllostomidae, increased body size predated dental specialisations.

Carnivory in phyllostomids is thought to have evolved twice convergently from a insectivorous common ancestor (Baker et al. 2012). The four carnivorous phyllostomid species are split between two tribes in the Phyllostominae: Phyllostomini (*Phyllostomus hastatus*) and Vampyrini (*C. auritus, T. cirrhosus* and *V. spectrum*) (Hoffmann et al. 2008, Wetterer et al. 2000; Baker et al. 2016), both of which also include non-carnivorous species. The convergent evolution of carnivory in Chiroptera has been studied at the ordinal level (evolving at least six times (e.g. Datzmann et al. 2010, Norberg and Fenton 1988, Santana and Cheung 2016), and the family level (evolving at least twice in Megadermatidae; e.g. Hand 1985, 1996). Independent evolutionary shifts to increasing body size in Vampryini and Phyllostomini from a smaller phyllostomid ancestor also support two instances of convergent evolution of carnivory (Arévalo 2020). Similarities in DTA between carnivorous species of both tribes in our study indicate they converged in the same morphospace.

### La Venta bat community

Authors of palaeoecological reconstructions have inferred the La Venta palaeoenvironment to have been a warm, wet and aseasonal mosaic woodland, possibly with patches of open grasslands and riverine systems (Croft 2001, Kay and Madden 1997, Spradley et al. 2019). Palaeogeographic models of the eastern Andean Cordillera suggest La Venta had a palaeoelevation of < 200 m, reflecting a lowland ecosystem (Guerrero 1997, Hoorn 1994, Hoorn et al. 1995). Precipitation has been estimated in the range of 1500-2000 mm and temperatures around 20-25 degrees Celsius, similar to the conditions found in modern forests of the Andes-Amazonian transition of tropical South America (Kay and Madden 1997, Spradley et al. 2019). This type of transitional ecosystem is characterised by rich taxonomic diversity and high endemism, possibly due to reciprocal faunal exchange between highland and lowland tropical faunas since the Miocene (Upham et al. 2013).

The fossil bat community is also informative for palaeohabitat reconstructions of La Venta and supports the presence of mosaic woodlands. Two modern species of foliage-roosting species (*Thyroptera* spp.) typical of lowland tropical forests have been reported from La Venta, indicating the presence of *Heliconia*-like vegetation (Czaplewski et al. 2003). The presence of the trawling insectivore *N. albiventris* in the La Venta fossil fauna/community also indicates the presence of bodies of water in the ecosystem (Kalko et al. 1998). Although this species is mainly an insectivore, some of its features (enlarged feet and constant frequency echolocation) have been interpreted as pre-adaptations for piscivory (Kalko et al. 1998), supported by different studies reporting the consumption of insects, fruits and pollen by *N. albiventris* (Fernando et al. 2007). Dietary reconstruction of La Venta’s *P. antimaster* indicates it was an omnivore that included nectar in its diet (Yohe et al. 2015). Our dietary reconstruction for *N. magdalenensis* and a previous reconstruction for *P. antimaster* (Yohe et al. 2015) corroborate the hypothesised mosaic woodland ecosystem of La Venta, with a mixture of high canopy trees and understory flowering vegetation (Kay and Madden 1997).

The 14 bat species recorded from La Venta would have exploited at least three types of food resources (insectivory, omnivory and nectarivory), occupying the trawling, gleaning and hawking aerial guilds. They occupied a wide body size range (4-50 g), implying an ecologically diverse community (Czaplewski et al. 2003). Our results indicate *N. magdalenensis* was the largest known noctilionoid bat in the La Venta fauna, although body mass reconstructions of the other extinct species would better elucidate the trophic structure of the bat community. The interpretation of *N. leporinus, P. antimaster* and *N. magdalenensis* as transitional forms (Paván et al. 2013, Yohe et al. 2015, Simmons et al. 2020) suggests ecosystems like La Venta may have been important for the evolution of dietary innovations (i.e. carnivory, insectivory and nectarivory) in Phyllostomidae (Kalko et al. 1998, Yohe et al. 2015). Further study of the La Venta bat fauna is expected to further inform hypotheses regarding the evolution of Phyllostomidae, the most ecologically diverse family of mammals.

## Conclusions

We used dental topography analysis and a quantitative comparative approach to reconstruct the diet of the phyllostomine *Notonycteris magdalenensis* from the middle Miocene La Venta fauna (tropical South America), the most diverse fossil bat community in South America. Our study investigated patterns of dental complexity (DNE, OPCR and RFI) variation across seven dietary guilds and provided the first multivariate DTA in Chiroptera. We found strong differentiation in dental complexity between dietary guilds, indicating that DTA is an informative tool to study ecomorphology in bats. Applying phylogenetic comparative methods, we found statistical support for the presence of both ecological and phylogenetic signal in the variation of molar complexity. Our results suggest *N. magdalenensis* was an omnivore or insectivore, rather than a carnivore like its extant sister taxa *C. auritus* and *V. spectrum*. Based on our BM reconstructions of *N. magdalenensis* (∼50 g), we infer it was the largest predatory bat in La Venta, being larger than most modern insectivorous bats, but smaller than most modern carnivorous bats. Combining our diet and BM reconstructions, we interpret *N. magdalenensis*’s BM and dental topography to represent pre-adaptations associated with the colonisation of a carnivorous niche. Our results confirm that *Notonycteris magdalenensis* was probably not a specialised carnivore. Evidence that modern carnivory (typically associated with a large body mass and specialised cranial morphology) had evolved in Phyllostomidae by the middle Miocene remains to be demonstrated.

## Acknowledgements

We would like to thank Tzong Hung and the facilities and technical assistance of the National Imaging Facility, a National Collaborative Research Infrastructure Strategy (NCRIS) capability, at the Mark Wainwright Analytical Centre, UNSW Sydney; and Sharlene Santana for granting access to the *V. spectrum* scans. CL-A is supported by a Research Training Program (RTP) scholarship from the Australian Department of Education and a Science PhD Writing Scholarship from the University of New South Wales, and SJH is supported by the Australian Research Council. Casts of UCMP specimens were provided by the late Donald E. Savage.

---

**Supplementary Table 1.** Data and references used in this study.

| Species | Diet | DNE | OPCR | RFI | BM | Diet reference |
| --- | --- | --- | --- | --- | --- | --- |
| *Anoura geoffroyi* | Nectarivore | 213.780 | 67.00 | 0.5203 | 15 | Arbour et al., 2019 |
| *Artibeus obscurus* | Frugivore | 407.930 | 128.12 | 0.4244 | 36.86 | Arbour et al., 2019 |
| *Carollia brevicauda* | Omnivore | 218.650 | 56.25 | 0.5832 | 14.33 | Santana et al., 2011 |
| *Chrotopterus aurita* | Carnivore | 281.590 | 102.50 | 0.5927 | 80.31 | Santana and Cheung, 2016 |
| *Desmodus rotundus* | Sanguivore | 124.656 | 28.38 | 0.7655 | 33.46 | Arbour et al., 2019 |
| *Diaemus youngi* | Sanguivore | 138.490 | 39.00 | 0.6466 | 36.59 | Arbour et al., 2019 |
| *Diphylla ecaudata* | Sanguivore | 92.263 | 17.50 | 0.6572 | 30.05 | Arbour et al., 2019 |
| *Glossophaga leachii* | Nectarivore | 165.120 | 52.50 | 0.4718 | 10.58 | Arbour et al., 2019 |
| *Leptonycteris curasoae* | Nectarivore | 177.180 | 35.00 | 0.5566 | 23.54 | Arbour et al., 2019 |
| *Lonchorhina aurita* | Insectivore | 244.640 | 64.88 | 0.5585 | 15.33 | Santana et al., 2011 |
| *Lophostoma silvicolum* | Insectivore | 556.688 | 179.12 | 0.5634 | 31.06 | Santana et al., 2011 |
| *Micronycteris schmidtorum* | Insectivore | 343.620 | 74.00 | 0.5185 | 7.63 | Santana et al., 2011 |
| *Mimon crenulatum* | Insectivore | 469.058 | 86.12 | 0.6183 | 13.56 | Santana et al., 2011 |
| *Musonycteris harrisoni* | Nectarivore | 170.480 | 52.50 | 0.5083 | 11.68 | Arbour et al., 2019 |
| *Noctilio albiventris* | Omnivore | 393.829 | 70.00 | 0.6274 | 30.03 | Fernando et al., 2007 |
| *Noctilio leporinus* | Piscivore | 509.362 | 85.25 | 0.7341 | 60.49 | Arbour et al., 2019 |
| *Phylloderma stenops* | Omnivore | 271.560 | 71.75 | 0.6054 | 53.84 | Santana et al., 2011 |
| *Phyllostomus elongatus* | Omnivore | 499.550 | 94.38 | 0.6536 | 40.37 | Santana et al., 2011 |
| *Phyllostomus hastatus* | Carnivore | 387.630 | 86.62 | 0.6018 | 93.52 | Santana and Cheung, 2016 |
| *Platyrrhinus vittatus* | Frugivore | 637.292 | 150.75 | 0.5095 | 62.3 | Arbour et al., 2019 |
| *Sturnira bogotensis* | Frugivore | 127.870 | 45.75 | 0.3500 | 20.51 | Arbour et al., 2019 |
| *Tonatia saurophila* | Insectivore | 291.350 | 97.38 | 0.6832 | 28.56 | Santana et al., 2011 |
| *Trachops cirrhosus* | Carnivore | 375.514 | 72.75 | 0.6333 | 35.63 | Santana and Cheung, 2016 |
| *Vampyressa pusilla* | Frugivore | 315.030 | 87.12 | 0.4807 | 9.08 | Arbour et al., 2019 |
| *Vampyrum spectrum* | Carnivore | 231.420 | 65.5 | 0.7411 | 177.29 | Santana and Cheung, 2016 |
| *Notonycteris magdalenensis* |  | 286.680 | 67.00 | 0.5688 | 53.05 |  |

**Supplementary Table 2.** Posterior probabilities of LDA including all three DTA metrics (DNE, OPCR and RFI).

| Species | Carnivore | Frugivore | Insectivore | Nectarivore | Omnivore | Piscivore | Sanguivore |
| --- | --- | --- | --- | --- | --- | --- | --- |
| *Anoura geoffroyi* | 0.071 | 0.057 | 0.192 | 0.592 | 0.086 | 0.000 | 0.002 |
| *Artibeus obscurus* | 0.000 | 0.978 | 0.011 | 0.010 | 0.000 | 0.000 | 0.000 |
| *Carollia brevicauda* | 0.254 | 0.003 | 0.177 | 0.261 | 0.268 | 0.002 | 0.035 |
| *Chrotopterus aurita* | 0.343 | 0.006 | 0.447 | 0.105 | 0.091 | 0.000 | 0.007 |
| *Desmodus rotundus* | 0.016 | 0.000 | 0.000 | 0.000 | 0.003 | 0.002 | 0.979 |
| *Daemus youngi* | 0.243 | 0.000 | 0.027 | 0.036 | 0.105 | 0.002 | 0.586 |
| *Diphylla ecaudata* | 0.087 | 0.000 | 0.005 | 0.013 | 0.049 | 0.002 | 0.844 |
| *Glossophaga leachii* | 0.012 | 0.130 | 0.057 | 0.774 | 0.026 | 0.000 | 0.000 |
| *Leptonycteris curasoae* | 0.121 | 0.004 | 0.093 | 0.491 | 0.262 | 0.001 | 0.027 |
| *Lonchorhina aurita* | 0.168 | 0.014 | 0.238 | 0.356 | 0.216 | 0.001 | 0.008 |
| *Lophostoma silviculum* | 0.038 | 0.199 | 0.742 | 0.007 | 0.013 | 0.000 | 0.000 |
| *Micronycteris schmidtorum* | 0.045 | 0.177 | 0.250 | 0.326 | 0.201 | 0.001 | 0.000 |
| *Mimon crenulatum* | 0.127 | 0.003 | 0.196 | 0.018 | 0.516 | 0.139 | 0.001 |
| *Musonycteris harrisoni* | 0.043 | 0.042 | 0.103 | 0.743 | 0.067 | 0.000 | 0.002 |
| *Noctilio albiventris* | 0.179 | 0.001 | 0.135 | 0.026 | 0.536 | 0.117 | 0.007 |
| *Noctilio leporinus* | 0.028 | 0.000 | 0.005 | 0.000 | 0.055 | 0.909 | 0.003 |
| *Phylloderma stenops* | 0.343 | 0.002 | 0.234 | 0.115 | 0.274 | 0.003 | 0.029 |
| *Phyllostomus elongatus* | 0.140 | 0.000 | 0.135 | 0.005 | 0.391 | 0.327 | 0.002 |
| *Phyllostomus hastatus* | 0.216 | 0.005 | 0.320 | 0.059 | 0.381 | 0.015 | 0.003 |
| *Platyrrhinus vittatus* | 0.002 | 0.838 | 0.141 | 0.005 | 0.014 | 0.000 | 0.000 |
| *Sturnira bogotensis* | 0.000 | 0.804 | 0.001 | 0.194 | 0.000 | 0.000 | 0.000 |
| *Tonatia saurophila* | 0.640 | 0.000 | 0.136 | 0.008 | 0.091 | 0.004 | 0.121 |
| *Trachops cirrhosus* | 0.242 | 0.001 | 0.154 | 0.027 | 0.484 | 0.079 | 0.013 |
| *Vampyressa pusilla* | 0.012 | 0.524 | 0.147 | 0.283 | 0.033 | 0.000 | 0.000 |
| *Vampyrum spectrum* | 0.203 | 0.000 | 0.007 | 0.001 | 0.031 | 0.014 | 0.745 |
| *Notonycteris magdalenensis* | 0.176 | 0.012 | 0.254 | 0.241 | 0.309 | 0.002 | 0.006 |

